# Resistify - A rapid and accurate annotation tool to identify NLRs and study their genomic organisation

**DOI:** 10.1101/2024.02.14.580321

**Authors:** Moray Smith, John T. Jones, Ingo Hein

## Abstract

**Background:** Nucleotide-binding domain Leucine-rich Repeat (NLR) proteins are a key component of the plant innate immune system. In plant genomes NLRs exhibit considerable presence/absence variation and sequence diversity. Recent advances in sequencing technologies have made the generation of high-quality novel plant genome assemblies considerably more straightforward. Accurately identifying NLRs from these genomes is a prerequisite for improving our understanding of NLRs and identifying potential novel sources of disease resistance.

**Results:** Whilst several tools have been developed to predict NLRs, they are hampered by low accuracy, speed, and availability. Here, the NLR annotation tool Resistify is presented. Resistify is an easy-to-use, rapid, and accurate tool to identify and classify NLRs from protein sequences. Applying Resistify to the RefPlantNLR database demonstrates that it can correctly identify NLRs from a diverse range of species. Applying Resistify in combination with tools to identify transposable elements to a panel of Solanaceae genomes reveals a previously undescribed association between NLRs and Helitron transposable elements.

**Conclusion:** Resistify can rapidly identify NLRs within plant genomes and provides accurate structural classifications. Its ease of use and accessibility allows easy integration into bioinformatic workflows and projects, enhancing the study of this important group of genes. Applying Resistify to a *Solanaceae* pangenome reveals an undescribed association between NLRs and transposable elements.

Availability: https://github.com/SwiftSeal/resistify

## Background

The plant innate immune system is an essential part of plant disease resistance. A key component of this are the Nucleotide-binding domain Leucine-rich Repeat (NLR) proteins – an abundant and diverse family present in plant and animal genomes. NLRs act as intracellular immune receptors which elicit an immune response after the detection of pathogen derived factors or their activity (Biezen and Jones, 1998). Since their discovery, hundreds of functional NLRs have been cloned and many sequences have been identified (Kourelis and van der Hoorn, 2018). The abundance of NLRs and their sequence diversity shows considerable variation across the plant kingdom (Barragan and Weigel, 2021).

One driving force of NLR diversification is thought to be the proliferation of transposable elements (TEs). In *Arabidopsis*, sites of high TE insertion frequency have been previously identified to contain NLRs (Quadrana *et al*., 2016), and NLRs exhibiting elevated intraspecific diversity are in close proximity to TEs (Sutherland *et al*., 2023). Of the TEs, LTR-retrotransposons are the most prolific in plant genomes and are a major factor in genome expansion (Lee and Kim, 2014). Strikingly, the *Capsicum annuum* genome exhibits a marked expansion of NLRs nested within LTR-retrotransposons, and this effect is seen in other *Solanaceae* including tomato and potato (Seong *et al*., 2020). Other transposable elements, such as the Helitrons, can also capture and transport genes across the plant genome (Lai *et al*., 2005).

Canonical NLRs have a modular organisation. Central to the NLR structure is the highly conserved NB-ARC domain which regulates the activity of the protein. Downstream of this is the Leucine-Rich-Repeat (LRR) domain which has a role in ligand binding and auto-inhibition. Upstream of the NB-ARC domain is variable. Often, there is either a coiled-coil (CC), Resistance to Powdery Mildew 8-like (RPW8), or Toll/Interleukin-1 receptor/Resistance protein (TIR) domain. Together, this modular structure means that NLRs can be separated into different classifications: CNL, RNL, TNL, or NL if lacking a conserved N-terminal domain.

Recently, additional NLR associated signatures have been identified. For example, the post-LRR C-terminal jelly-roll/Ig-like domain (C-JID) (Martin *et al*., 2020) may be present in TNLs and appears to bind with pathogen effectors, thereby contributing to the immune response. Similarly, other NLRs have integrated decoys to detect pathogen activities. In terms of initiating cell death through the formation of a resistosome, CNLs rely on an N-terminal MADA motif, a hallmark of NRCs - NLRs which act as intermediaries in the NLR induced immune response or NLRs that function independently of NRCs (Adachi *et al*., 2019).

Due to their conservation and modular structure, NLRs lend themselves to automated identification and classification. To date, several tools have been developed to achieve this including DRAGO2, NLGenomeSweeper, NLR-Annotator, RGAugury, RRGPredictor, and NLRtracker (table 1) (Li *et al*., 2016; Osuna-Cruz *et al*., 2018; Santana Silva and Micheli, 2020; Steuernagel *et al*., 2020; Toda *et al*., 2020; Kourelis *et al*., 2021). The development of the RefPlantNLR database has allowed the benchmarking of these tools against the sequences of functionally characterised NLRs (Kourelis *et al*., 2021). Performance varied between tools with NLRtracker being the most sensitive and accurate. All tools perform well at the identification of TIR, RPW8, and NB-ARC domains which are highly conserved, but are often less accurate in predicting CC domains which are more variable and frequently missed by InterProScan (Jones *et al*., 2014). Additional to the currently available NLR classification tools is NLRexpress – a set of machine learning predictors for CC, TIR, NB-ARC, and LRR motifs (Martin *et al*., 2022). Although NLRexpress does not directly identify or classify NLRs, it is well suited for rapidly and accurately screening large sets of sequences for NLR-associated motifs. One strength of NLRexpress is the CC-associated extended EDVID model which performs strongly in identifying this challenging motif.

**Table 1:** A summary of the currently available NLR annotation tools. Adapted from (Kourelis et al., 2021).

| Tool | Input data | Method | Output | Distribution |
| --- | --- | --- | --- | --- |
| **DRAGO2** | Protein,<br>Transcript | Custom HMMs | Classification,<br>Domains | Online only,<br>API available |
| **NLGenomeSweeper** | Transcript,<br>Genomic | InterProScan,<br>MUSCLE,<br>TransDecoder,<br>BLAST,<br>HMMER | Classification,<br>Genome<br>position,<br>GFF annotation | GitHub,<br>Manual<br>dependency<br>installation |
| **NLR-Annotator** | Transcript,<br>Genomic | MEME motifs | Classification,<br>Genome<br>position,<br>GFF annotation | GitHub |
| **RGAugury** | Transcript,<br>Genomic | InterProScan | Classification, | GitHub, |
|  |  |  | Genome<br>position,<br>GFF annotation | Online or local<br>webservice,<br>Docker<br>container |
| **RRGPredictor** | Protein,<br>Transcript | InterProScan | Classification | GitHub,<br>Manual<br>dependency<br>installation |
| **NLRtracker** | Protein,<br>Transcript | InterProScan,<br>HMMER,<br>MEME motifs | Classification,<br>NB-ARC<br>sequence,<br>Domains,<br>GFF annotation | GitHub,<br>Manual<br>dependency<br>installation |
| **NLRexpress** | Protein | HMMER,<br>Scikit-learn | Motif position,<br>Motif<br>sequence | GitHub,<br>Online or local,<br>Conda<br>environment<br>provided |

One drawback with the currently available tools is the reliance on InterProScan as the backend for domain annotation. InterProScan is designed as a comprehensive and generalised domain annotation tool. As a result, each input sequence is scanned against several databases each containing in total more than 180,000 protein signatures, the vast majority of which are not NLR associated. Compounding this, NLRs only represent a fraction of a plant proteome resulting in unnecessary searches against non-NLR sequences. In addition, domains common to NLRs are present in InterProScan databases with differing levels of curation which often results in overlapping or fragmented annotations. These must be parsed, particularly the LRR domain which is represented by multiple InterProScan signatures. The large databases that InterProScan relies upon must also be downloaded which provides an additional challenge for installation or automated deployment.

Software that is readily available with minimum manual configuration is a requirement for genome assembly and annotation projects. Projects may use an array of different tools, making manual installation and dependency resolution time-consuming. Dependency resolution solutions such as Conda, and automated deployment of tools in pipelines with Snakemake or Nextflow are becoming increasingly common (Di Tommaso *et al*., 2017; Mölder *et al*., 2021). Additionally, bioinformatics research is often restricted to shared High-Performance Clusters (HPCs) where users have limited privileges and where storage is at a premium. Tools that require root privileges, are distributed as web services, or require large databases are often not conducive for genomics projects.

Here, the new NLR annotation tool Resistify is presented that overcomes some of the constraints of currently available tools. We show that this tool accurately predicts NLR sequences from diverse plant using a range of well curated sources and can be used for pangenome analysis of NLRs in *Solanaceous* genomes. In addition, we use Resistify in combination with the EDTA tool to investigate the genomic organisation of NLRs and their association with transposable elements.

## Methods

Resistify is implemented in python as a command line executable. First, Resistify executes a hmmsearch of input protein sequences against a custom database of HMMs derived from curated Pfam entries (table 2). These models are used to identify CC, RPW8, TIR, and NB-ARC domains, as well as C-JID and MADA motifs. Domains of the same type are merged if they overlap or are within 100 amino acids of each other. Preliminary testing in development showed that this is necessary to overcome NB-ARC domain annotations which can become split. MADA motifs are restricted to the N-terminal of the sequence. Using this evidence, proteins are initially classified as belonging to either a CN, RN, TN, or N classification. Proteins which do not have any evidence of an NB-ARC domain are discarded at this stage.

**Table 2:**
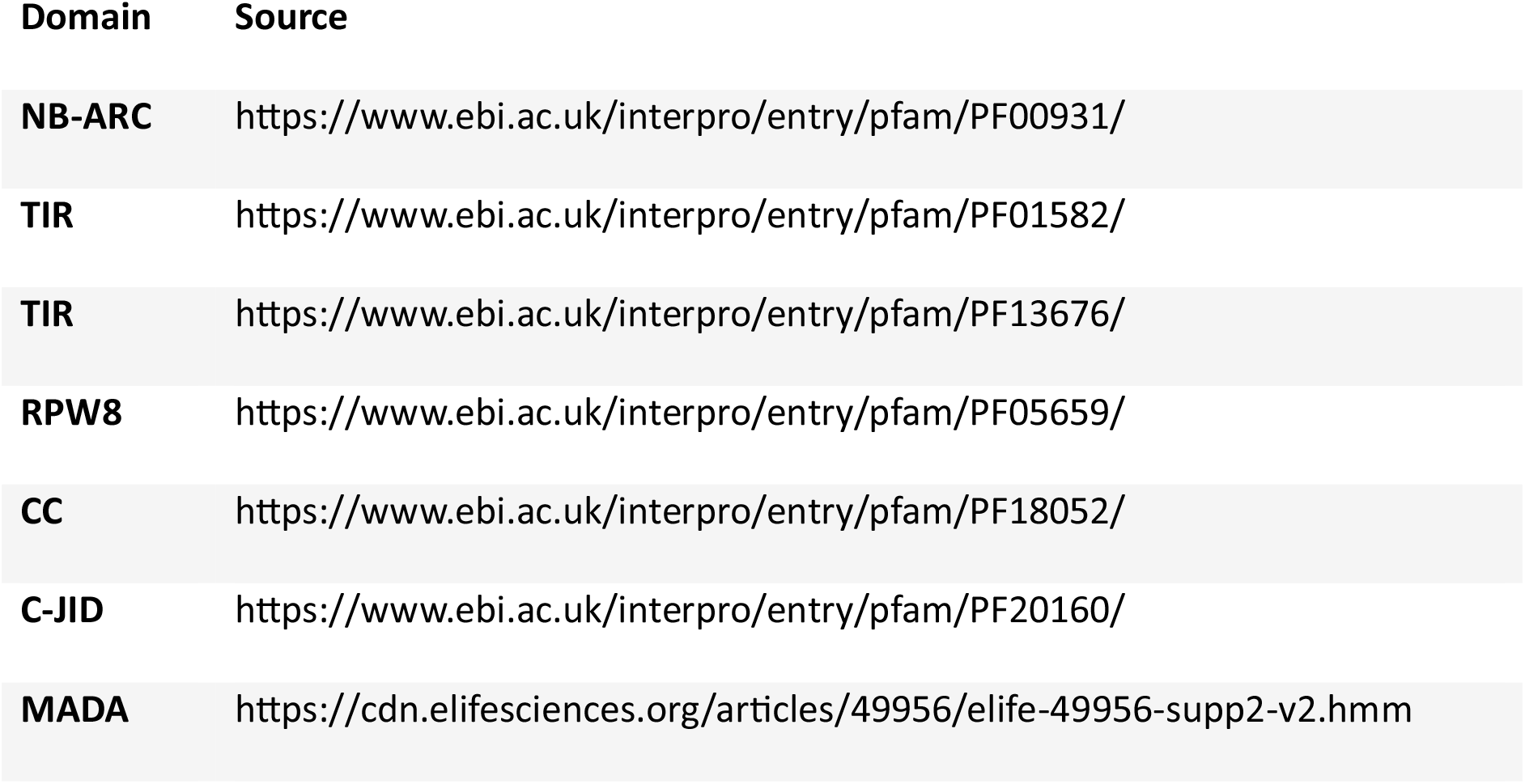
HMM models including in the initial hmmsearch stage of Resistify.

Following this, the filtered set of NB-ARC containing proteins is screened for NLR-associated motifs with the NLRexpress models. Sequences are then reclassified using the following logic. If a protein belongs to class N (i.e., does not have any evidence of an upstream CC, RPW8, or TIR domain) it is scanned for upstream TIR or CC motifs. As motif searches can be more promiscuous, restricting motif searches to this condition prevents them from interfering with non-ambiguous NLR classification. The sequence is then screened for LxLxxL motifs to define the LRR domain. Following a previous definition, an LRR domain is annotated if four or more LxLxxL motifs are identified with inter-motif gaps of less than 75 amino acids [@martin_nlrscape_2023]. Gaps larger than 75 amino acids are predicted to be a break in the LRR domain and so the LRR annotation process is restarted from that position onwards. If less than four motifs exist across the whole sequence this process is skipped.

This combined evidence is then integrated into the domain annotation data. The domains are sorted by start position and a “domain string” is formed. For example, if the sorted domains took the order TIR, NB-ARC, LRR, C-JID, then the domain string would be TNLj. Alternatively, a canonical CNL with a MADA motif would take the form mCNL. The domain strings are searched for substrings CNL, RNL, TNL, or NL and classified accordingly. Sequences are further classified on the basis of whether they contain an N-terminal bound MADA motif or CJID domain. Additionally, a count of the unique occurrences of the conserved NB-ARC motifs is made as an indicator of NB-ARC integrity.

The primary output of Resistify is a table detailing the NLRs identified and the specific classification for each sequence. The complete domain string, classification, NB-ARC motif count, and MADA and CJID status are listed. A complete list of all annotations and NLRexpress motifs are given as separate tables. Additionally, all NB-ARC domain sequences are extracted and provided as a FASTA file.

Resistify is provided as a single executable available for download on GitHub, PyPi, and Conda. All databases and models are distributed with the executable so that manual setup is not required. All output files are placed in a nameable output directory for easy integration into automated workflows. Temporary files are also handled internally to reduce output clutter. Resistify requires scikit-learn v0.24.2 and hmmer v>=3.0.

### RefPlantNLR benchmarking

Protein sequences of the RefPlantNLR database members were retrieved and used as input for Resistify with default settings (Kourelis *et al*., 2021). Resistify classifications were compared directly in R v4.3.2 with tidyverse v2.0.0. Any sequence where the Resistify classification did not exactly match with the provided RefPlantNLR structure was taken as a potential misclassification.

### Araport11 benchmarking

The latest release of Araport11 representative gene model protein sequences were downloaded from TAIR and used as input for Resistify with default settings(Lamesch *et al*., 2012). A phylogenetic tree was built from the Resistify-extracted NB-ARC domain sequences with mafft v7.52.0 and fasttree v2.1.11 with default settings.

### Pangenome pipeline

A Snakemake workflow was developed to predict genes and NLRs in a Solanum pangenome comprised of chromosome-scale genomes. Snakemake was executed in a mamba v1.4.2 environment with snakemake v7.32.3 and cookiecutter v1.7.3 (Mölder *et al*., 2021). Genes were predicted *de-novo* from the genome alone using Helixer v0.3.2 with the land_plant_v0.3_a_0080.h5 model (Holst *et al*., 2023). Protein sequences were extracted using AGAT v1.2.0 and used as input for Resistify (Dainat *et al*., 2023). Ortholog analysis was carried out using OrthoFinder v2.5.5 using all predicted protein sequences (Emms and Kelly, 2019). Transposable elements were annotated with the latest GitHub release of EDTA (Ou *et al*., 2019).

Gene models were translated to bed format using AGAT v1.2.0 and overlaps with intact transposable elements were identified using bedtools v2.31.1 with the command bedtools intersect using the options -f 0.9 -wo (Quinlan and Hall, 2010). To identify previously characterised NLRs, protein sequences of a subset of *Solanum*-originating NLRs in the RefPlantNLR database were queried against each genome with blastp v2.15.0 (Camacho *et al*., 2009). The full pipeline and all post-hoc analysis are available at https://github.com/SwiftSeal/pangenomics/.

## Results

### Performance against RefPlantNLR

To evaluate the performance of Resistify (figure 1), it was applied against the RefPlantNLR database - a curated set of 415 previously cloned NLRs from a diverse range of species.

**Figure 1:**
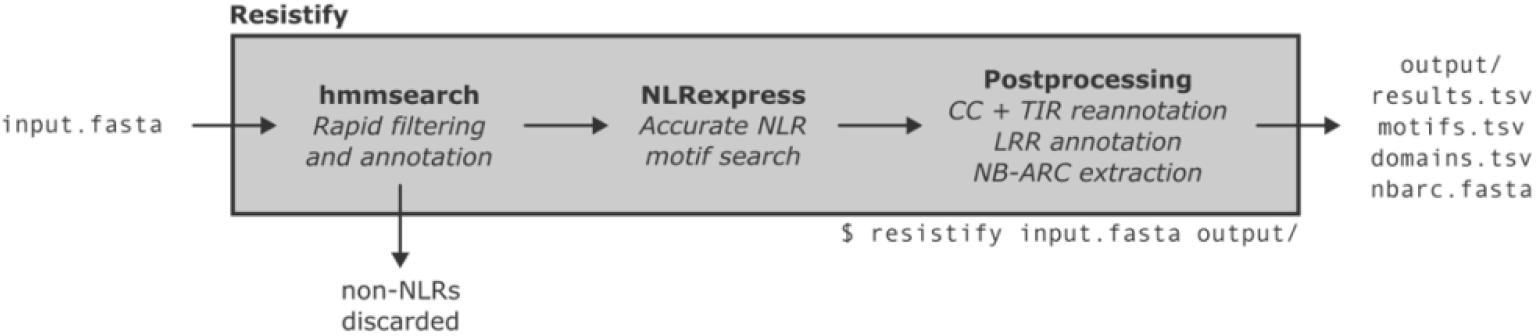
The Resistify program.

NLRtracker was also applied to the RefPlantNLR database for comparison.

In the default mode, only three RefPlantNLR entries were not identified by Resistify - *AtNRG1.3* which carries only an LRR domain and *Pb1* and *RXL* which have significantly truncated NB-ARC domain that are not listed in the RefPlantNLR database. However, if required, these genes can be identified with Resistify using –ultra mode which skips the initial filtering stage and reports and sequence which contains at least one NLRexpress motif. Sequences are reported as an unmerged string of NLR motifs. Consequently, *AtNRG1.3*, *Pb1*, and *RXL* are reported as NNLLLLLLL, CNNNLLLLLLLLLLL, and CNNNLLLLLLLLL respectively.

The largest source of variation between RefPlantNLR and Resistify classification was 29 NLRs which had an NL structure according to RefPlantNLR but CNL according to Resistify. All of these belonged to CNL-associated subclasses which are known to have CC domains that are challenging to identify. This included 15 CC_G10_-NLRs including *Pvr4*, *Tsw*, *RPS2*, *RPS5*, *SUT1* and *SUMM2* which have previously been noted for their lack of the CC-associated EDVID motif (Lee *et al*., 2021). Others included six members of the *Pm5* locus which despite not containing a CC domain in RefPlantNLR, have previously been identified to contain CC-like domains (Xie *et al*., 2020). This analysis demonstrates that Resistify is highly sensitive at retrieving canonical NLRs and accurately describing their structure.

As NLRtracker does not exclude NLRs without NB-ARC domains, *AtNRG1.3*, *Pb1* and *RXL* were identified and classified as “CC-NLR or CCR-NLR or CCG10-NLR”, “CC-NLR”, and “CC-NLR” respectively. NLRtracker relies on the NB-ARC associated RNBS-D motif to classify NLRs as “CC-NLR or CCR-NLR or CCG10-NLR”. As a result, 26 of the conflicting CNLs identified by Resistify were classified as “CC-NLR or CCR-NLR or CCG10-NLR” despite failing to identify a CC domain. The exception of this was for *SpNBS-LRR*, *Rpp1-R1*, and *Rpp4C4* which were classified as “UNDETERMINED”.

Reliance on RNBS-D also led to classification of *NtTPN1* as “CC-NLR or CCR-NLR or CCG10-NLR”. Whilst *NtTPN1* is a CNL ortholog, it lacks any upstream domain and is structurally an NL. Unexpectedly, the TNL *DSC1* was misclassified as “CC-NLR” despite having a domain structure of “(TIR)(NBARC)(LRR)(CJID)” according to NLRtracker.

In summary, Resistify performs well at identifying canonical NLRs from a diverse range of species. In default mode, it does not assign genes as NLRs with extremely truncated or entirely absent NB-ARC domains, unlike NLRtracker which reports any sequence with NLR-associated domains. However, this can be replicated in Resistify with the –ultra mode. Thus, Resistify’s structure-based classification method is well suited for correctly classifying NLRs, including members of challenging CNL subclasses.

### Performance against the Araport11 proteome

To assess the performance of Resistify across a well characterised and annotated genome, the representative gene models of Araport11 were analysed (Cheng *et al*., 2017). In total, Resistify identified 166 NLRs - the majority of which were TNLs and CNLs (figure 2a). Of the CNLs, 25% had a MADA motif, and 44.4% and 41.2% of NLs and TNLs had C-JID domain respectively. Partial NLRs either without an N-terminal or LRR domain were also identified.

**Figure 2:**
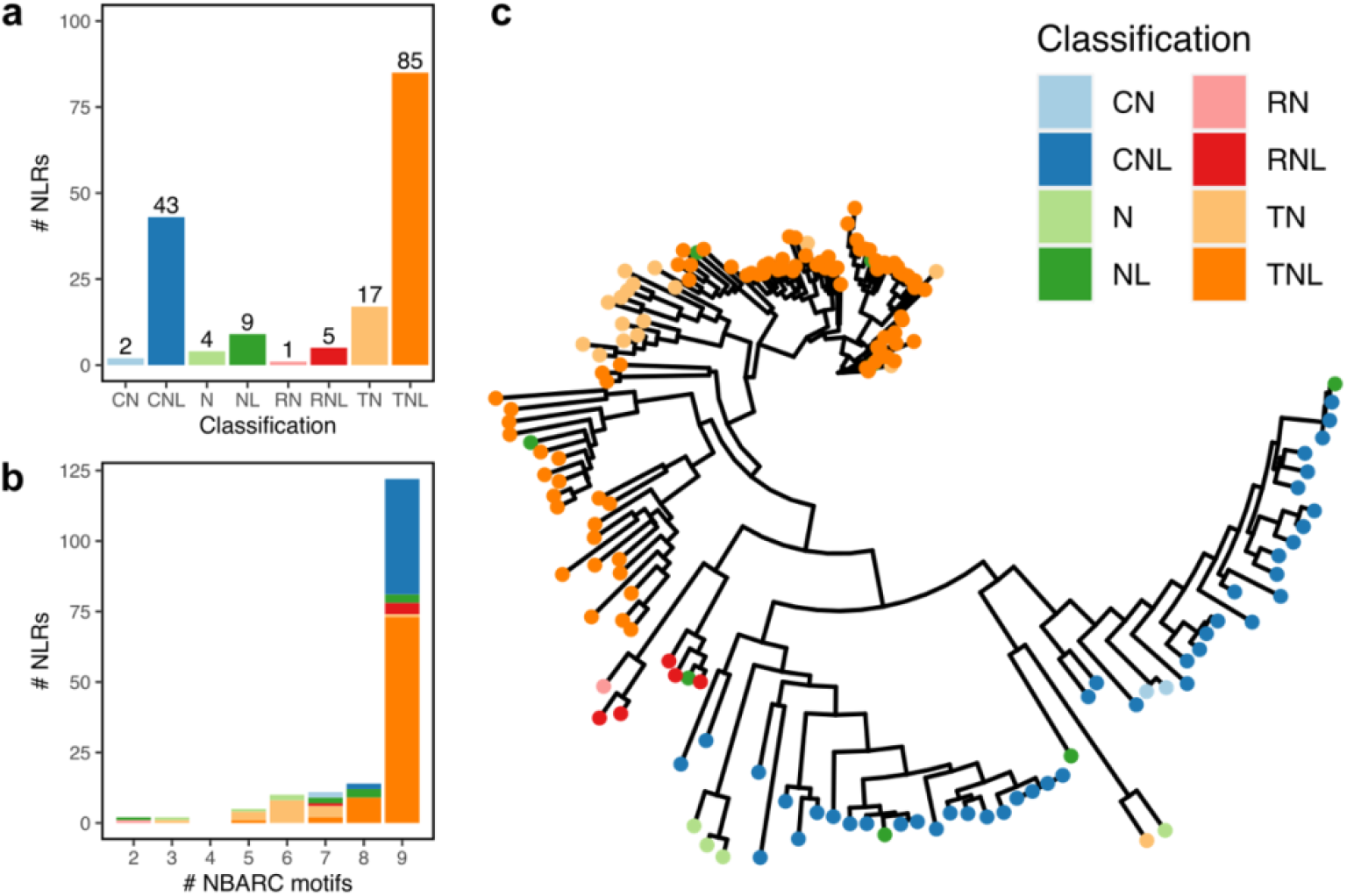
Resistify applied to Araport11. a) The number of NLRs belonging to each identified classification. b) The number of NLRs grouped by number of identified unique NB-ARC associated motifs. c) A phylogenetic tree of NLRs based on the Resistify extracted NB-ARC domain sequences.

Manual inspection of the motifs within these sequences confirmed that they were not due to a failure to identify these elements. The majority of NLRs carried all nine conserved NB-ARC motifs - those with fewer increasingly belonged to partial NLR classifications (figure 2b). NLRs with as few as two of the conserved NB-ARC motifs were successfully identified. A phylogenetic tree of the Resistify-identified NB-ARC domains independently validated the classifications and allowed the placement of ambiguous NL or N classified sequences into subclasses (figure 2c).

NLRtracker identified an additional 48 sequences, however these sequences did not contain any NB-ARC domain annotation according to either tool. Three sequences - AT4G19060.1, AT4G19060.1, and AT5G45440.1 - were not identified by NLRtracker. According to Resistify these contained a single NB-ARC domain each, with 5, 6, and 6 NB-ARC motifs respectively. Overall, Resistify performs well at identifying and classifying NLRs from whole proteomes and successfully retrieves NB-ARC domain annotations for phylogenetic analyses.

### Application against an example workflow

To demonstrate how Resistify might be implemented to identify novel resistance genes, a pangenome experiment was performed. Eighteen contiguous *Solanum* genomes were downloaded and processed with a simple workflow which predicts genes de novo, identifies orthologues, and classifies NLRs with Resistify (Tang *et al*., 2022; Li *et al*., 2023). For gene annotations, the recently developed tool Helixer was selected for its near reference quality predictions and lack of requirement for repeat-masking which is known to result in false-negative NLR annotations (Bayer, Edwards and Batley, 2018; Holst *et al*., 2023). The workflow also predicts transposable elements with EDTA. The *C. annuum* genome was included as an outgroup, but also because recent analysis has suggested a substantial expansion of NLRs in this species associated with LTR transposable elements (Kim *et al*., 2017).

Predicted gene content varied between genomes from 29,223 in *S. habrochaites* to 61,015 in the larger *C. annuum* genome (table S1). Transposable element content ranged from 55.6% in *S. chmielewskii* to 76.8% in *C. annuum*. Whilst there was no significant difference in total TE content between tuber-bearing and non-tuber-bearing genomes (p = 0.644), there was a significant increase in the number of intact TEs reported by EDTA in the genomes of tuber-bearing species (p = 0.008).

In total, 8144 NLRs were identified across all genomes (figure 3). CNLs were the most abundant classification of NLR identified, ranging from 84 in *S. habrochaites* to 422 in *S. tuberosum* (group tuberosum RH10-15). There was a notable expansion of NLRs in tuber-bearing *Solanum* species. This is in agreement with the previous observation that potato-bearing *Solanum* species have an expansion of tuber expressed NLRs (Tang *et al*., 2022). This effect does not correlate with the proportion of genome occupied by transposable elements of any classification (figure S1).

**Figure 3:**
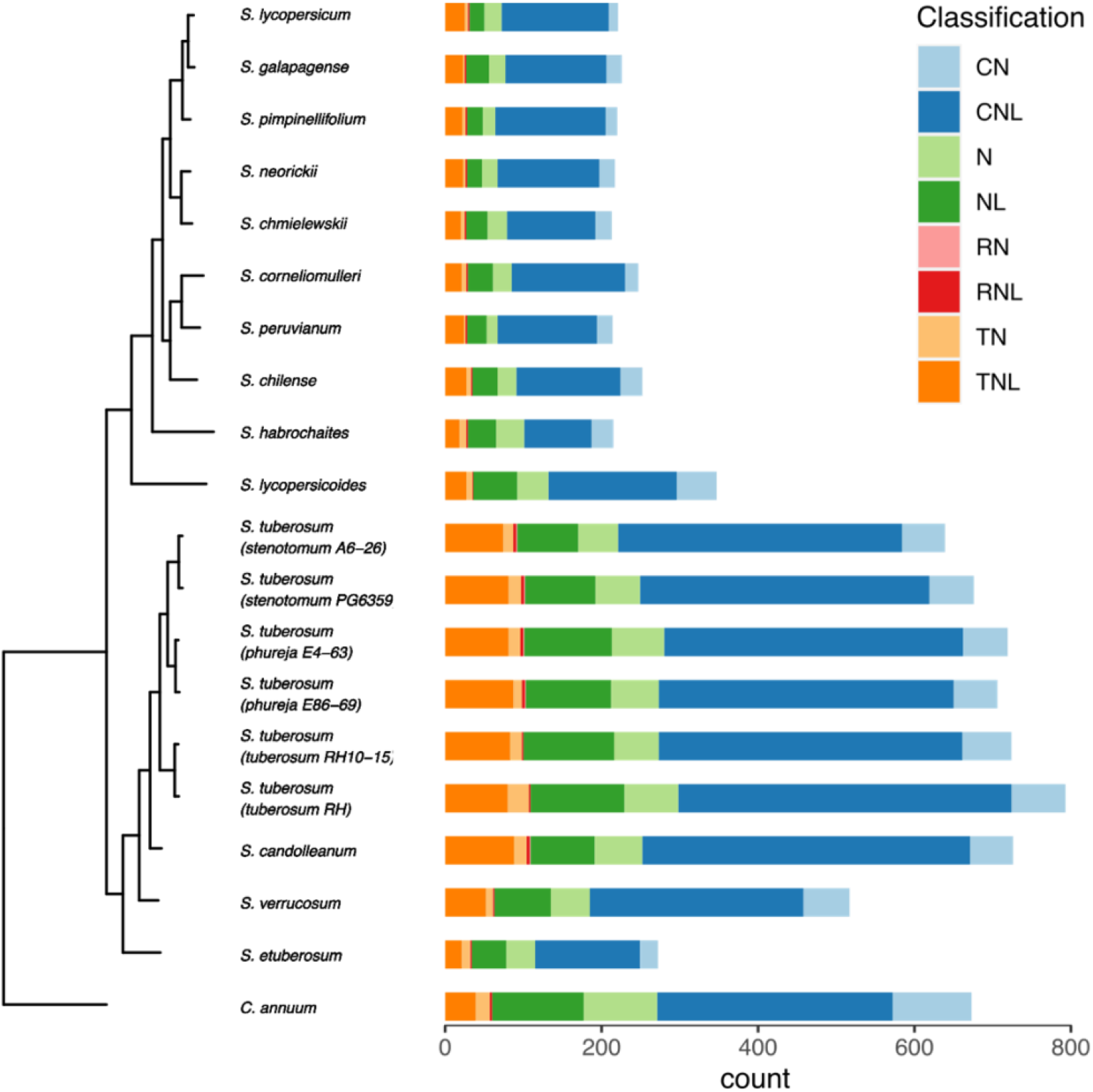
NLRs identified across the *Solanum* pangenome.

In total, 38,590 orthogroups were identified, of which 687 (1.8%) contained NLRs. The distribution of orthogroups showed a clear divide between core and shell/cloud orthogroups within the pangenome with the majority of NLR orthogroups existing within the shell/cloud (fig 4a). This reflects the relatively large width of the pangenome, which captures NLR variation over a Genus level. Species specific NLRomes also exhibit an abundance of cloud orthographs, reflecting the high variability of NLRs in genomes (Van de Weyer *et al*., 2019). Many (48.1%, n = 371) orthogroups were classified as containing N or NL NLRs (figure 4b). Whilst these classifications are less abundant (24.6%, n = 2081), they were often associated with smaller orthogroups contributing to this inflation.

**Figure 4:**
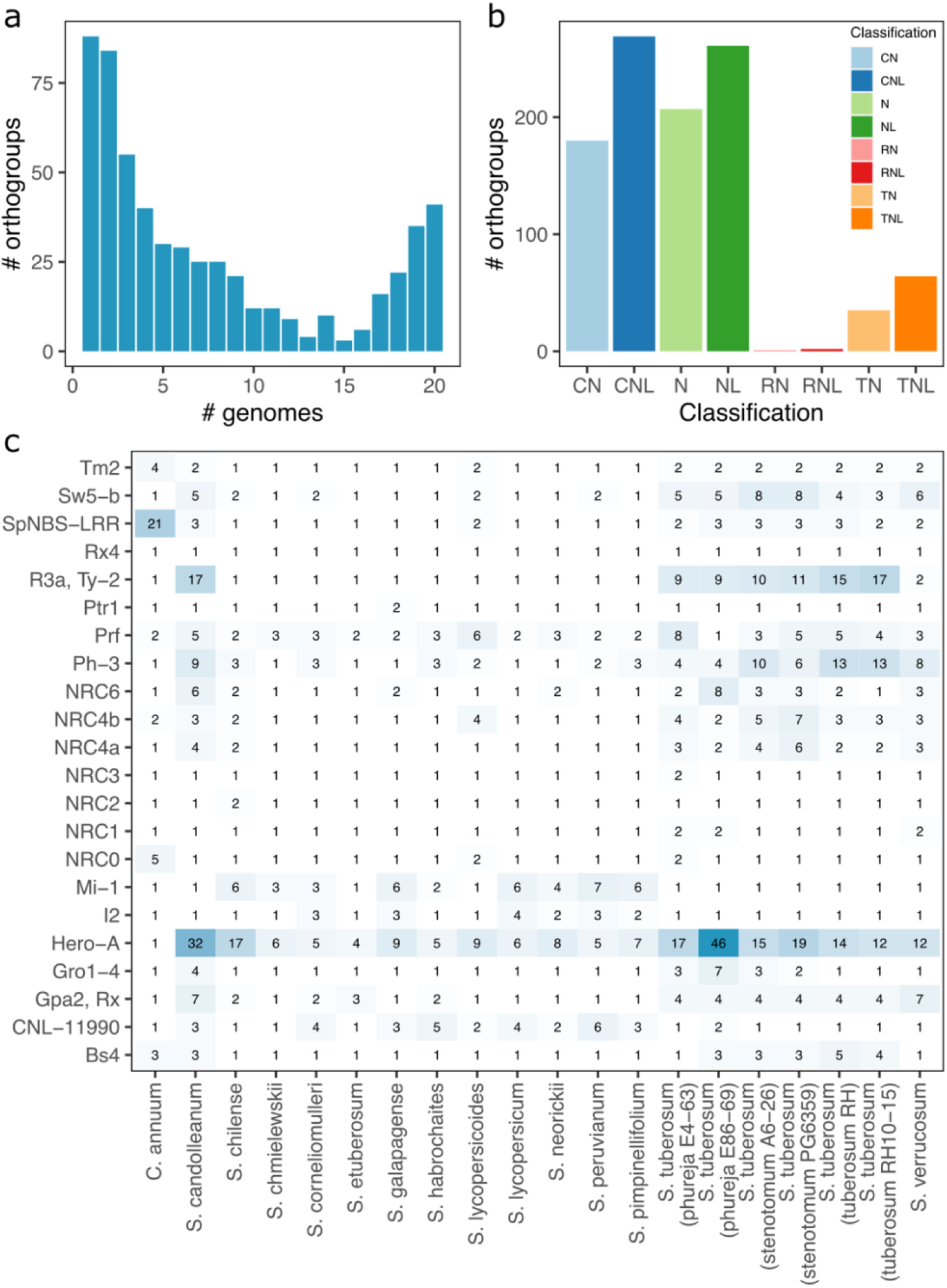
NLR-containing orthogroups. a) The number of orthogroups shared between genomes. b) The number of orthogroups according to NLR classification. c) The number of orthologs of known *Solanaceae* NLRs identified in each genome.

The distribution of previously identified *Solanaceae* NLR orthologs across the pangenome was examined (figure 4c). In agreement with previous findings, members of the NRC group remained relatively stable across the pangenome except for NRC4a, NRC4b, and NRC6 which were expanded in the tuber-bearing genomes. By contrast, an expansion of *Mi-1* and *Hero* was evident in the non-tuber bearing and tuber bearing genomes respectively. At least one ortholog was identified in each genome for all genomes.

It has previously been reported that there is a vast expansion of NLRs in the *C. annum* genome due to retroduplication; NLRs nested within LTRs represented a large proportion (∼13%) of NLRs within the genome, and this effect is also seen in tomato (8%) and potato (18%) (Kim *et al*., 2017). To explore whether this effect could be linked to the expansion of NLRs within the tuberising members of *Solanum*, a similar analysis was repeated. Intact TEs were identified and considered to interact with NLRs if they covered >90% of the gene annotation.

Unexpectedly, the effect could not be replicated and across all genomes only five intact LTRs were identified as containing NLRs (figure 5a). None were identified in *C. annuum* but instead in the *S. tuberosum* group Phureja and clone RH. The putative retrotransposed NLRs within these all belonged to the same orthogroup (OG0000639) which was expanded in both Phureja and RH, but not *S. stenotomum*.

**Figure 5:**
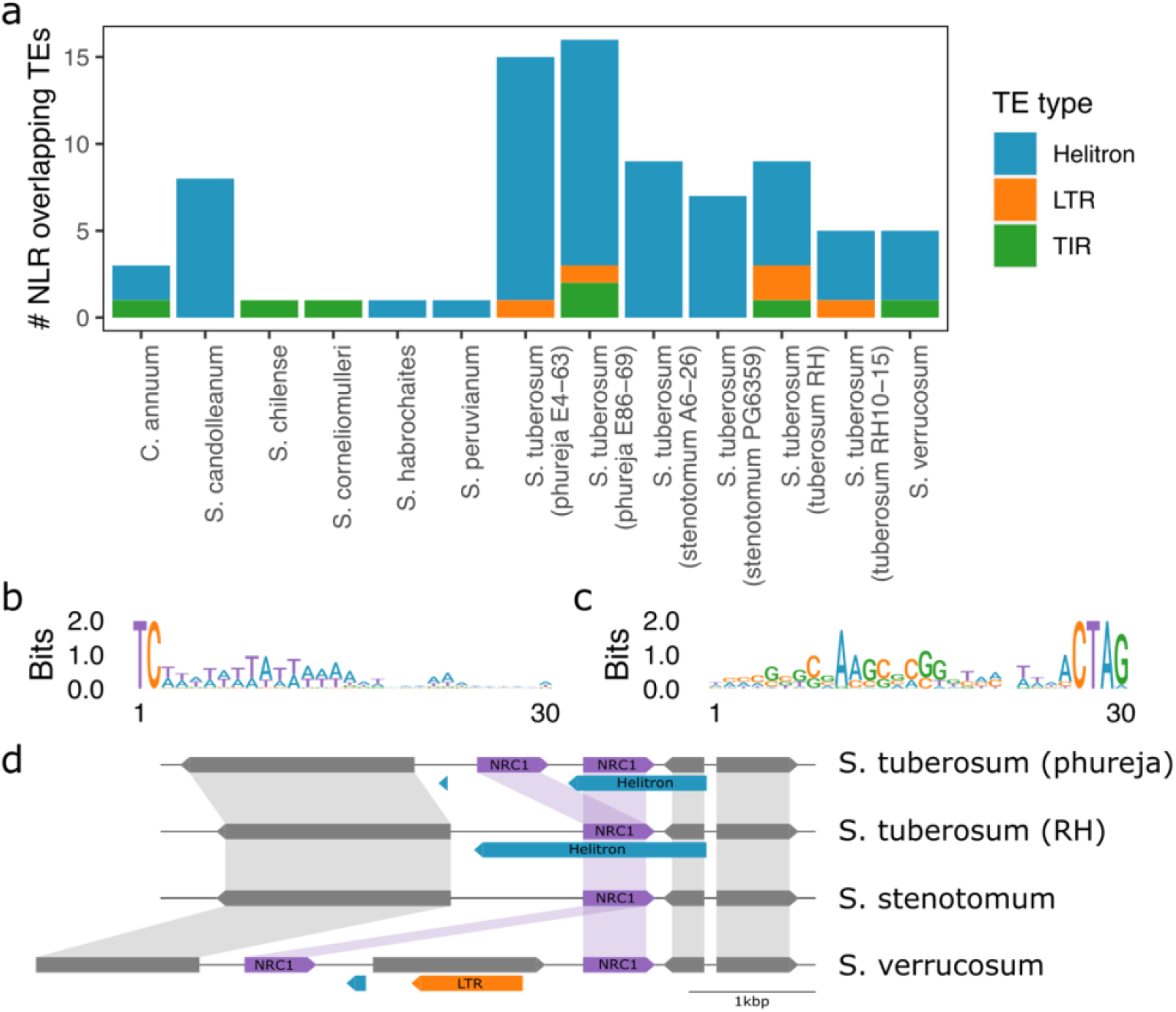
Helitrons associated with NLRs. a) The number of intact TEs overlapping with NLRs across the pangenome. b) The 5’ motif of NLR overlapping Helitrons. c) The 3’ motif of NLR overlapping Helitrons. d) The *NRC1* locus and its association with Helitrons across four genomes. Helitron (blue) and LTR (orange) transposable elements are highlighted.

By contrast, a proliferation of Helitron-associated NLRs was identified in the tuber-bearing genomes. All predicted Helitrons carried the expected 5’-TC…CTRR-3’ signature as well as a GC-rich region in the 3’ terminal (figure 5b,c). An example of an NLR which may have undergone Helitron-associated duplication is *NRC1* (figure 5d) which is present as a single copy except in both *S. tuberosum* group Phureja genomes and *S. verrucosum*. Close examination of the *NRC1* locus reveals a duplication event in *S. tuberosum* group Phureja where one *NRC1* gene is nested entirely within a predicted Helitron. In *S. tuberosum* clone RH which only has one copy of *NRC1*, *NRC1* is similarly nested within a Helitron although this appears to be extended to a distant Helitron terminator past the site of duplication in group Phureja. In *S. stenotomum*, only one *NRC1* is present and no Helitron predicted.

Interestingly, *S. verrucosum* has two copies of *NRC1* and whilst it does not have a nested Helitron copy, the leftmost *NRC1* copy does have a short predicted Helitron upstream of its start position in the same orientation as the other Helitron elements. In this case, the two copies of *NRC1* appear to have been further separated by an intact LTR insertion which has resulted in an additional gene prediction.

## Discussion

Resistify is presented as a highly accurate and easy-to-use tool to aid the identification of novel resistance genes. Applying Resistify against the RefPlantNLR database demonstrates that it is effective at identifying NLRs from a diverse range of plant genomes. It is highly sensitive at classifying challenging CNL families which often evade other NLR classification tools due to the inclusion of the performant NLRexpress motif models. Members of the CC_G10_-NLR clade have previously been described as lacking the CC-associated EDVID motif and are often predicted as having an NL structure in the RefPlantNLR database (Lee *et al*., 2021). By utilising NLRexpress’ highly sensitive extended EDVID motif model, Resistify successfully classifies these elusive NLR clades, highlighting it as the most sensitive NLR classifier to date.

Resistify can be easily integrated into workflows and is scalable to large pangenome experiments. As high-quality genome assemblies become more common, providing accessible tools for the study of NLRs will be crucial for fully appreciating their diversity and identifying novel sources of resistance. The recent releases of chromosome level assemblies for potato and tomato pangenomes are a valuable resource for understanding the diversity of NLRs within *Solanum* (Tang *et al*., 2022; Li *et al*., 2023). An expansion of NLRs including essential families such as the NRCs is apparent in tuber-bearing species.

As Resistify relies on an initial NB-ARC domain search to reduce the search space prior to NLRexpress motif identification, NLRs with very truncated or entirely absent NB-ARC domains can be missed. To resolve this, an additional mode is provided which does not perform initial NB-ARC domain filtering which identifies sequences with any NLR motifs.

Unlike NLRtracker, Resistify does not search for noncanonical integrated domains (Kroj *et al*., 2016). As integrated domains are widespread and often critical to NLR function, their identification and analysis are an important factor in studies of NLR diversity (Barragan and Weigel, 2021). Pairing Resistify with a secondary sweep for these domains with NLRtracker or InterProScan would permit this analysis whilst also benefiting from a reduced search space and high-quality NLR classifications from Resistify.

An unexpected finding of this study was the failure to replicate the previous observation of an abundance of LTR associated NLRs across *Solanaceae* (Kim *et al*., 2017). Differences in gene and TE annotation and NLR classification likely contributed to this. Although Helixer predicted ∼70% more genes in *C. annuum* than were identified in the previous study, this did not translate to an increase in NLRs identified. Instead, only 673 NLRs were identified here in comparison to the previous estimate of 835. Previously, a tBLASTn search of the NB-ARC domain followed by ORF identification and BLASTP searches against GenBank to classify NLRs was used to identify NLRs in the genome (Seo *et al*., 2016). This method may result in more false positives in comparison to identifying NLRs directly from gene annotations. Here, EDTA was selected for identifying TEs which has an improved sensitivity and selectivity for identifying intact LTRs in comparison to LTRHarvest which was used previously. Although Helixer does not require repeat masking which can result in false negative NLR annotations, annotations used in training the model (which may have relied upon repeat masking) might introduce TE-avoidant behaviour (Bayer, Edwards and Batley, 2018). However, the identification of several LTR-associated NLRs indicates that this is unlikely to be the source of the difference in the result.

The identification of Helitron-associated NLR expansion has not been previously reported. Helitrons are challenging to identify due to their lack of structural elements and as a result the pipeline used in this study is known to overestimate Helitron density (Baril, Imrie and Hayward, 2022). However, a large part of this overestimation is likely due to EDTA reannotating the genome with HelitronScanner predictions, which itself suffers from a high false positive rate, leading to a proliferation of fragmented Helitron annotations across the genome. For this study, only intact Helitrons which passed EDTAs stringent filter were considered. As a result, all predicted Helitrons had the required structural elements for activity. Further validation would be required to determine if Helitron association is a valid mechanism of NLR expansion. The Helitron/*NRC1* relationship highlighted here would be a good starting point for unpicking this mechanism.

## Conclusions

Resistify is presented as a new tool to aide in the study of NLRs in plant genomes. It provides high-quality structural classifications for canonical NLRs. Applying Resistify to the RefPlantNLR database demonstrates that it can correctly classify NLRs from a diverse range of plant species. Applying Resistify to a panel of *Solanaceae* genomes demonstrates its suitability for studying NLRs in large datasets. This analysis revealed functionally important NLR families undergoing expansion in tuberising *Solanum* species. Unexpectedly, combining this with transposable element annotations did not identify a previous description of NLRs embedded within LTR retrotransposons in *Solanaceae* genomes. Instead, an undescribed association between NLRs and Helitron transposable elements in genomes with expanded NLR inventories was revealed.

## Acknowledgements

The authors acknowledge the Research/Scientific Computing teams at The James Hutton Institute and NIAB for providing computational resources and technical support for the “UK’s Crop Diversity Bioinformatics HPC” (BBSRC grant BB/S019669/1), use of which has contributed to the results reported within this paper. The authors also acknowledge Peter Cock for his support with creating the conda distribution of Resistify.

## Availability of data and materials

Resistify is available at https://github.com/SwiftSeal/resistify. All data analysis workflows and scripts are available at https://github.com/SwiftSeal/pangenomics. The RefPlantNLR database was accessed from its source publication (Kourelis *et al*., 2021). The genomes used in this study were accessed from their source publications (Tang *et al*., 2022; Li *et al*., 2023).

