## Supplementary figures and images for "Resistify - A rapid and accurate annotation tool to identify NLRs and study their genomic organisation"

### figure S1

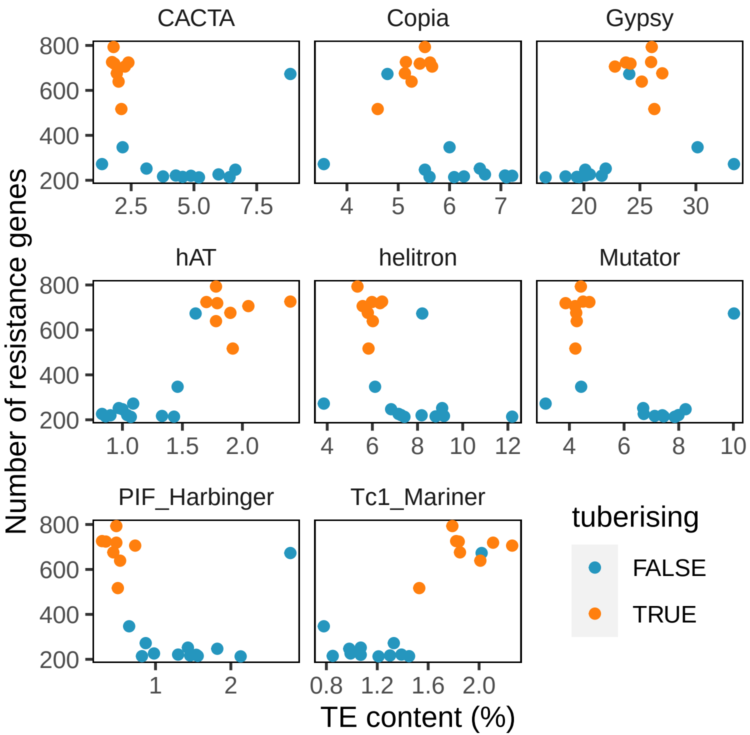
